# Prediction of RNA-protein sequence and structure binding preferences using deep convolutional and recurrent neural networks

**DOI:** 10.1101/146175

**Authors:** Xiaoyong Pan, Peter Rijnbeek, Junchi Yan, Hong-Bin Shen

## Abstract

RNA regulation is significantly dependent on its binding protein partner, which is known as the RNA-binding proteins (RBPs). Unfortunately, the binding preferences for most RBPs are still not well characterized, especially on the structure point of view. Informative signals hiding and interdependencies between sequence and structure specificities are two challenging problems for both predicting RBP binding sites and accurate sequence and structure motifs mining.

In this study, we propose a deep learning-based method, iDeepS, to simultaneously identify the binding sequence and structure motifs from RNA sequences using convolutional neural networks (CNNs) and a bidirectional long short term memory network (BLSTM). We first perform one-hot encoding for both the sequence and predicted secondary structure, which are appropriate for subsequent convolution operations. To reveal the hidden binding knowledge from the observations, the CNNs are applied to learn the abstract motif features. Considering the close relationship between sequences and predicted structures, we use the BLSTM to capture the long range dependencies between binding sequence and structure motifs identified by the CNNs. Finally, the learned weighted representations are fed into a classification layer to predict the RBP binding sites. We evaluated iDeepS on verified RBP binding sites derived from large-scale representative CLIP-seq datasets, and the results demonstrate that iDeepS can reliably predict the RBP binding sites on RNAs, and outperforms the state-of-the-art methods. An important advantage is that iDeepS is able to automatically extract both binding sequence and structure motifs, which will improve our transparent understanding of the mechanisms of binding specificities of RBPs. iDeepS is available at https://github.com/xypan1232/iDeepS.

## 1 Introduction

RNA-binding proteins (RBPs) are highly involved in various regulatory processes, e.g. gene splicing and localization [7]. Understanding the roles of individual RBPs in gene regulation requires the detailed characterization of the binding preferences, which are short regions on RNAs, also called sequence motifs. Currently, there are many high-throughput technologies, e.g. RIP-seq and CLIP-seq, to determine binding sequence motifs of individual RBPs. However, they are time-intensive and costing.

The available high-throughput data of a large number of reliable RBP binding sites can serve as a gold standard for training and testing of less-expensive and faster prediction models [26, 1]. Many sequence-motif discovery tools have been developed. For example, the widely used MEME model fits a mixture model using expectation maximization to discover multiple sequence motifs [2]. MatrixREDUCE infers the sequence-specific binding motifs for transcription factors [8].

However, pervious studies have shown that many RBPs bind to RNA molecules by recognizing specific sequences, but also secondary structure contexts [14, 24]. The sequential motifs alone cannot fully explain the binding preference of the RBPs. For instance, the amyotrophic lateral sclerosis associated protein FET binds to its RNA target within hairpin loops structure [16].

Considering the impact of structure context on RBP binding preferences, some tools have been developed to identify binding RBP binding sites by taking also secondary structure into consideration. For instance, MEMERIS searches for RNA motifs enriched in regions with high structural accessibility [14]. Li et al., integrate the accessibility of RNA regions around the RBP interaction sites to identify accessible sequence motifs [24]. CapR models the joint distribution of residue positions and secondary structures to identify the binding sites under different structure context [9]. RNAcontext trains machine learning models using sequence and accessibility information to infer sequence and structure motifs [18]. GraphProt [26] integrates the RNA sequence and secondary structural contexts using a graph kernel model to investigate the RBP binding preferences, and it represents input sequences using over 30,000 dimensional graph features. Recently, the iONMF [35] integrates kmer sequence, secondary structure, CLIP co-binding, Gene Ontology (GO) information and region type using orthogonal matrix factorization to predict binding sites. However, the above methods require domain knowledge to construct the input features.

Alternatively, deep learning methods are fully data-driven. They automatically learns high-level features from simple input features, it simultaneously does feature engineering and model learning. Recently deep learning proved to be very successful in many research areas, e.g image recognitions [21, 13]. Also, promising performances were demonstrated on predicting RNA-protein interactions and binding sites [1, 29, 37]. For instance, DeepBind applies CNNs to automatically capture the binding sequence motifs [1]. Furthermore, our previous iDeep model predicts the RBP binding sites on RNAs and sequence motifs using the hybrid CNNs and DBNs through integrating multiple sources of representations [28]. However, similar to DeepBind [1], it can discover only the sequence binding preferences. Deepnet-rbp incorporates structure features into predicting the binding sites using deep belief networks (DBNs). It includes the RNA structure information, obtained from another tool, as a count vector of k-mers [37]. A disadvantage of Deepnet-rbp is that it requires complicate steps to estimate the binding preference.

In this study, we propose and evaluate a novel deep learning method, called iDeepS, which consists of CNNs and a bidirectional LSTM. This method identifies the sequence and structure binding motifs simultaneously. To the best of our knowledge, iDeepS is the first CNN-based study to easily capture both the sequence and structure binding motifs.

## 2 MATERIALS AND METHODS

We design the computational approach iDeepS (Figure 1) to predict the RBP binding sites on RNAs. We use sequences of RBPs binding sites derived from CLIP-seq dataset, apply one-hot encoding for the sequences and predicted secondary structures, and feed these into CNNs and a BLSTM to predict RBP binding sites. Finally, we extract the sequence and structure motifs from the learned convolution filters of the CNNs and evaluate them against known verified motifs.

**Figure 1:**
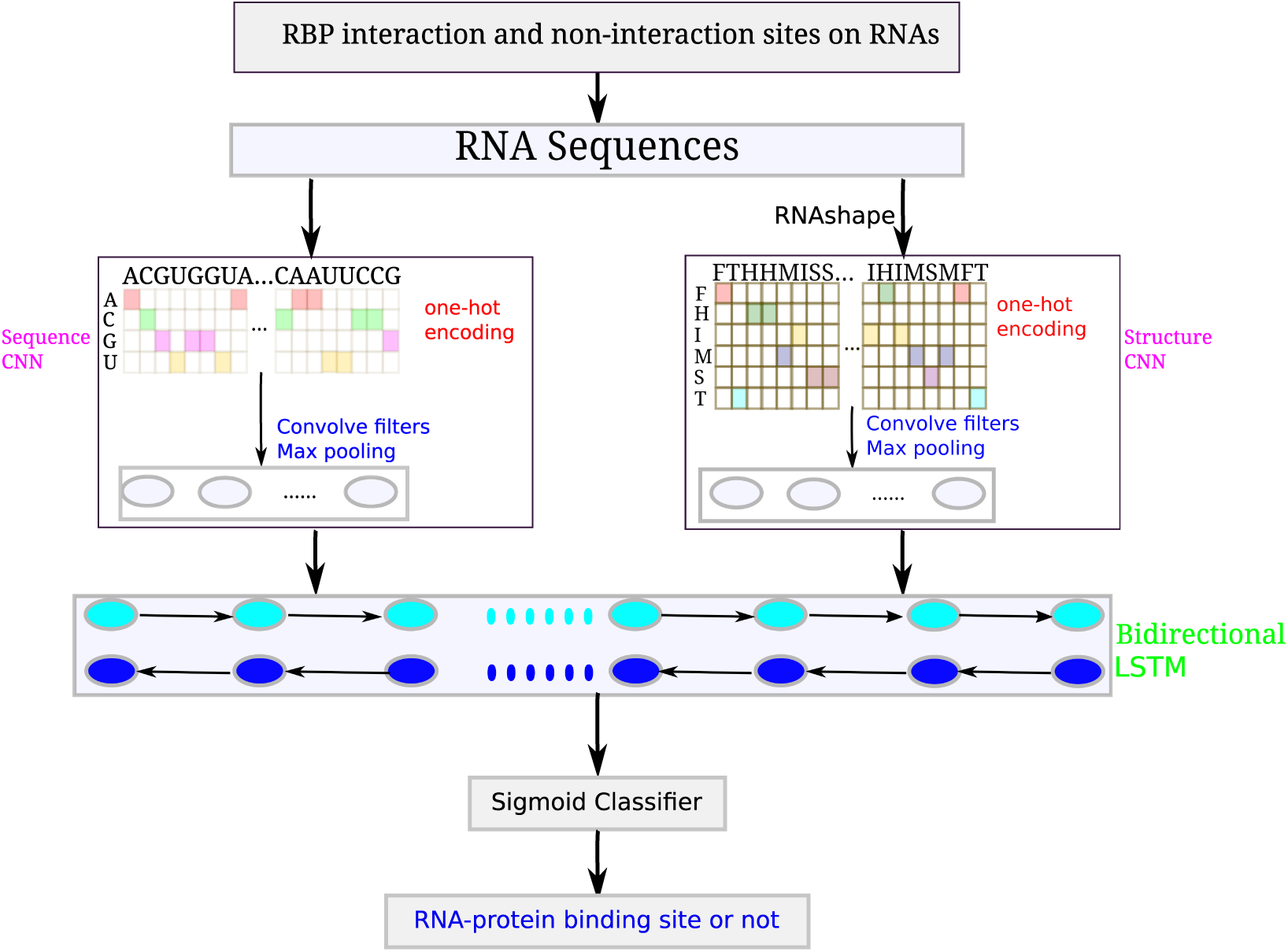
The flowchart of iDeepS. For each experiment, iDeepS integrates two CNNs (one is for sequences, the other is for structures predicted by RNAshape from sequences) to predict RBP interaction sites and identify binding sequence and structure motifs, followed by the bidirectional LSTM, which learns the long range dependencies between learned sequence and structure motifs. Finally, the outputs from bidirectional LSTM are fed into a sigmoid classifier to predict the probability of being RBP binding sites

### 2.1 Datasets

In this study, we train deep learning models for RBP binding sites derived from CLIP-seq data [35] available at (https://github.com/mstrazar/ionmf). This CLIP-seq dataset consists of 19 proteins with 31 experiments. For each experiment, each nucleotide within clusters of interaction sites derived from CLIP-seq were considered as binding sites. The negative sites were sampled from genes that were not identified as interaction in any of 31 experiments. In each experiment, a total 24,000 samples are used for training, 6,000 samples for model optimization and validation, and the other 1,0000 samples for independent testing.

### 2.2 Encoding sequence and structure

The RNA sequence is used as an one-hot representation encoded into a binary matrix, whose columns correspond to the presence of A, C, G, U and N [1, 38]. Given a RNA sequence *s* = (*s*_1_*, s*_2_*,…, s_n_*) with n nucleotides and sequence motif detector with defined size m, the binary matrix M for this sequence is represented as follows:
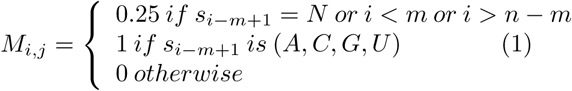

We use abstract secondary structure annotation from RNAshapes [34] implemented in https://github.com/fabriziocosta/EDeN. The RNAshapes have six generic shapes: stems (S), multiloops (M), hairpins (H), internal loops (I), dangling end (T) and dangling start (F). For each sequence s, we obtain the structure shapes *str* = (*str*_1_*, str*_2_*,…, str_n_*) by RNAshapes, which are converted into a binary matrix R with columns corresponding to the presence of F, H, I, M, S, T, and, and with k representing the predefined structure motif size.
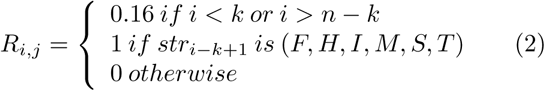

### 2.3 Convolutional neural network

The Convolutional Neural Network (CNN) [20] is inspired by the animal visual cortex. It consists of convolution, activation, and max-pool layers.

The one-hot encoding matrix derived from RNA sequences and structures are the inputs to the CNNs and are used to learn the weight parameters of the convolution filters. The convolution layer outputs the matrix inner product between input matrix and filters. After convolution, a rectified linear ReLU is applied to sparsify the output of the convolution layer and keep only positive matches to avoid the vanishing gradient problem [27]. Finally, a max pooling operation is used to reduce the dimensionality and yield invariance to small sequence shifts by pooling adjacent positions within a small window.

Before feeding into the next layer, the CNNs of sequence and structure are merged into one layer. The subsequent layers of the iDeepS act jointly on the merged sequence and structure layers.

### 2.4 Long Short Term Memory networks

LSTM belongs to the class of recurrent neural network [15], it incorporates long-term dependent information to assist the present prediction. In this study, LSTM is used to identify informative combinations of the extracted sequence and structure motifs [30], which projects the original input into a weighted representation.

As the LSTM sweeps across each element of the input, it first decides which information should be excluded by a forget gate layer based on previous inputs. Then an input gate layer is used to determine which information should be stored for next layer, and update the current state value. Finally, an output gate layer determines what parts of state value should output. Taking a sequence 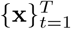 as input, the LSTM have the hidden states 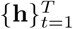, cell state 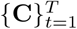, and it outputs a sequence 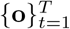. The above steps can be formulated as follows:

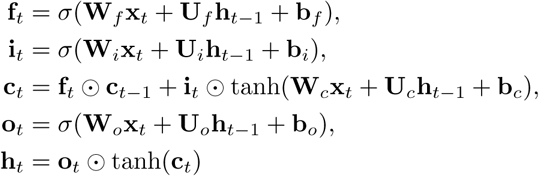

where ⊙ denotes element-wise multiplication, the *σ* is the Logistic Sigmod function and *tanh* is the tanh function to push the values to be between -1 and 1. **W***_f_*, **W***_i_*, **W***_o_*, **U***_f_*, **U***_i_* and **U***_o_* the weights and **b***_f_*, **b***_i_*, **b***_c_* and **b***_o_* the bias.

In iDeepS, bidirectional LSTM (BLSTM) is used, i.e., it sweeps from both left to right and right to left, and the outputs of individual directions are concatenated for subsequent classification.

### 2.5 Identifying the binding sequence and structure motifs

To explore the learned motifs, we investigate the convolve filters of sequence and structure CNNs in iDeepS. We convert them into position weight matrices (PWM) [1, 19], which are matched against input sequences and structures to discover binding motifs.

Assuming we have a sequence or structure *S_m_* and a convolve filter with size L, if the activation value *A_mfi_* of filter f at position i is greater than 0.5 max_*mi*_ A_*mfi*_, then this sequence or structure in windows L centring the position i is selected to align sequence motifs using WebLogo [6].

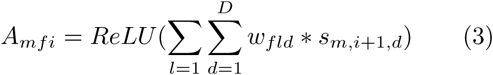

where *ReLU*(*x*) = *max*(0, *x*), *w_f_* is the weights of filter f, m is the length. For sequence motifs, D is 4. For structure motifs, D is 6.

To verify the predicted sequence motifs, we align them against 102 known motifs in study [31] from CISBP-RNA using the TOMTOM algorithm [10] with p-value <0.05. For some proteins, currently there are still no verified motifs in the CISBP-RNA database, we investigate them via the literatures.

Furthermore, we also calculate motif enrichment scores of predicted sequence and structure motifs using AME in the MEME suite [2]. Take sequence motifs as an example, it first scans the predicted motifs against the input sequences, and do the same for the shuffled sequences as the background sequences. Then we compare them to calculate the enrichment scores. We do the same enrichment analysis for predicted structure motifs.

### 2.6 Implementation

The iDeepS is implemented in python using keras 1.1.2 library https://github.com/fchollet/keras. We set the maximum number of epochs to 30, and the batch size to 50. The validation dataset is used to monitor the convergence during each epoch of the training process, so the training process can be stopped early. The model is trained by back-propagation using categorical cross-entropy loss, which is minimized by RMSprop [36]. In addition, we also employ multiple techniques to prevent or reduce over-fitting, e.g. batch normalization [17], dropout [33] and early stopping.

The number of motifs for both sequence and structure CNNs is set to 16 as suggested by DeepBind [1]. As indicated in iDeep [28], ReLU leads to information loss for some bits in motifs. As proposed by DeepBind, the the filter_length (motif width) should be 1.5 times the verified motif width, which is 7 in CISBP-RNA database [31]. Therefore, we choose a filter length of 10 in this study. When converting the filters to PWMs, we only use the first 7 bits of 10.

### 2.7 Baseline methods

There are many computational methods developed for predicting RNA-protein binding sites [1, 25, 26, 35]. In this study, we compare iDeepS with the state-of-the-art sequence-based methods DeepBind [1], Oli [25], iONMF [35] and Graph-Prot [26]. DeepBind, uses a sequence CNN with the same architecture as iDeepS to predict RBP binding sites. For GraphProt (v1.1.3), it encodes the sequence and structure into high-dimensional graph features, which are fed into a SVC to classify RBP bound and unbound sites. In this study, we use a window size 80 in GraphpProt and the other parameters are set to the default. iONMF uses matrix factorization to predict RBP binding sites by integrating different sources of features [35]. Oli uses linear SVC to classify RBP binding sites based on tetranucleotide frequency features [25]. The performance is measured using the area under the receiver operating characteristic curve (AUC).

## 3 RESULTS

In this study, we evaluate iDeepS on large-scale RBP binding sites from CLIP-seq. We evaluate the performance of iDeepS for predicting binding sites on RNAs, and compare it with the state-of-the-art methods. Furthermore, we identify the binding sequence and structure motifs using CNNs integrated in iDeepS.

### 3.1 Performance of iDeepS

To demonstrate the advantage of iDeepS, we compare it with the sequence-based DeepBind and Oli across the 31 experiments. iDeepS results in an average AUC of 0.86, which is a little better than 0.85 of DeepBind. The performance of Oli [25]is much lower than iDeepS, with an average AUC 0.77 across 31 experiments, which performs worse with a big margin than iDeepS (AUC 0.86). For some proteins, Oli’s performance is close to random guessing, e.g. protein hnRNPL with AUC 0.39. As showed in Figure 2 (More details are in Supplementary Table S1), iDeepS outperforms DeepBind on 25 of 31 experiments, and Oli on all experiments. It is interesting to note that the three methods have large performance differences across individual experiments. For iDeepS, the AUCs ranges from 0.59 for protein Ago2-MNASE to 0.98 for protein HNRNPC. For Ago2 protein, iDeepS cannot yield high performance. The reason is that Ago2 binding specificity is primarily mediated by miRNAs [3], the expressed miRNAs have an high influences Ago2-RNA interactions, which results in a more variable binding motifs than RBPs which bind RNAs directly.

**Figure 2:**
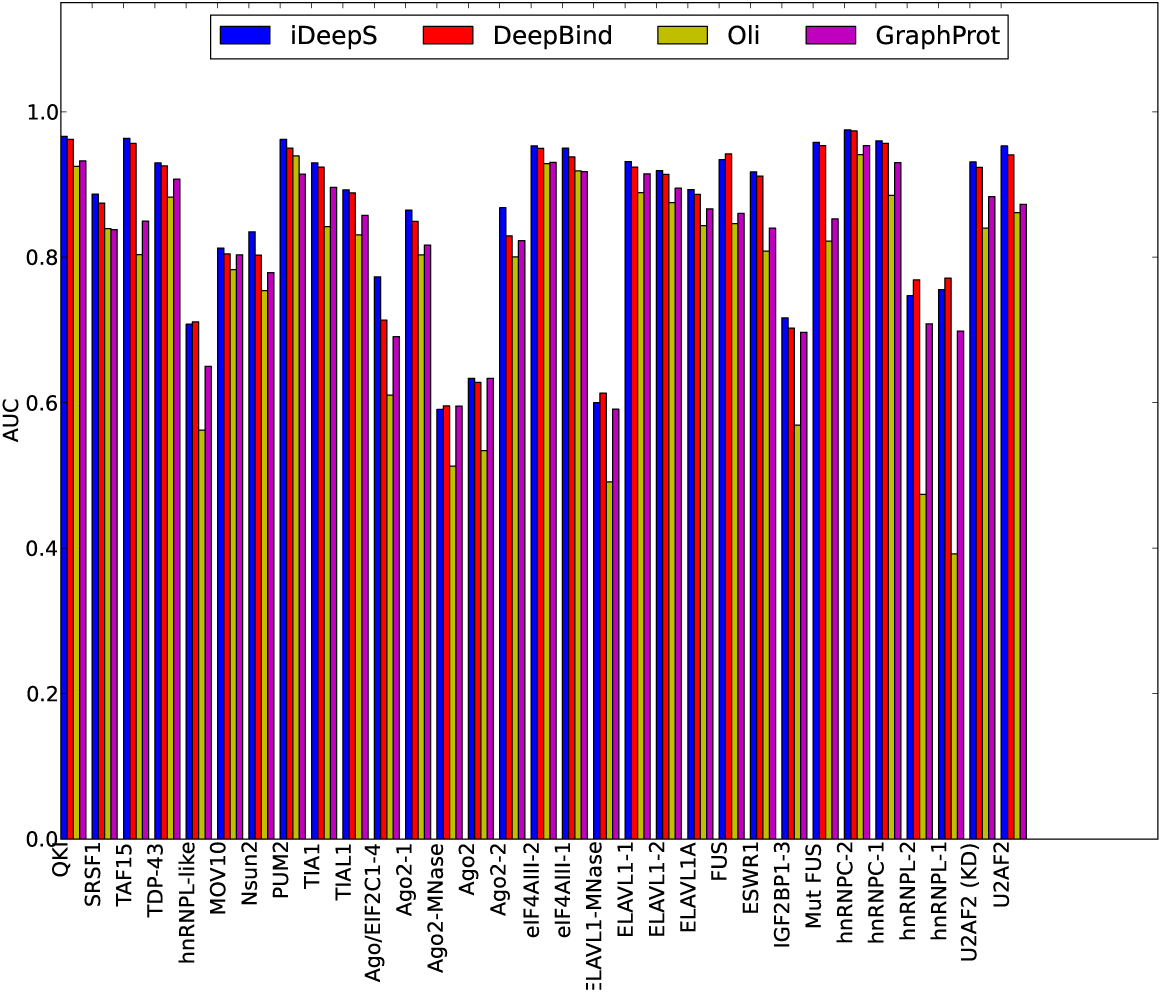
The AUCs of iDeepS, DeepBind, Oli and GraphProt across 31 experiments. The performances are evaluated on the same training and independent testing set across 31 experiments (x-axis) for iDeepS,DeepBind, Oli and GraphProt. For Oli and DeepBind, only sequences are used. For iDeepS and GraphProt, sequences and predicted structures are used.

We also compare iDeepS with structure-profile-based GraphProt, which demonstrates better performance than RNAcontext [26]. Across the 31 experiments, GraphProt yields the average AUC of 0.82, which is worse than 0.86 of iDeepS. As shown in Figure 2, iDeeps achieves better AUCs than GraphProt on 30 of 31 experiments. Our method improves the AUCs for some proteins with large margin. For example, iDeepS yields the AUC 0.77 for protein Ago/EIF, which is an increase of 12% compared to AUC 0.69 of GraphProt (Supplementary Table S1).

In addition, iDeepS outperforms iONMF (reported average AUC 0.85 on the same data) using multiple sources of data, including k-mer frequency, secondary structure, GO Information and gene type [35]. They also report that the iONMF surpasses the GraphProt and RNAcontext. The results further indicate that iDeepS can outperform other methods integrating other sources of hand-engineered features not derived from structures and sequences.

In summary, iDeepS not only on average achieves better performance than other peer sequence-based methods, it also outperforms other approaches integrating multiple sources of features. Our results demonstrate that iDeepS benefits strongly from learning the combination of high-level sequence and structure features for predicting binding sites for RBPs.

### 3.2 Insights in sequence-structure motifs

A big advantage of iDeepS is that it also provides the biological insights, e.g. learned binding motifs, of the RBPs by investigating the learned models. As compared to GraphProt, which requires a complicate postprocessing step, iDeepS easily converts learned parameters of the convolved filters and allows for identification of the sequence and structure motifs.

In this study, we infer the binding motifs across 31 experiments. Of them, 19 experiments have known sequence motifs in CISBP-RNA database or literatures. As shown in Table 1, iDeepS is able to discover experimentally verified sequence motifs for the 19 experiments. Of them, 15 are matched against CISBP-RNA with significant E-value cutoff 0.05 provided by TOMTOM [10]. The remaining 4 proteins resembles the motifs reported by other studies, which are based on the visual inspection. iDeepS discovers repeated UG dinucleotides motifs for TDP-43, which contains this dinucleotide repeats in 80% of the 3’UTR region by microarray analysis [26, 4], and captures a known motif, which plays a crucial regulator in germline development [32], for QKI with significant E-value 0.00008. The motif for PUM2 has been found with AU-rich sequence motif by iDeepS, which is close to the motifs identified based on top sequence read clusters [12]. The results show that the identified sequence motifs by iDeepS are aligned with verified motifs.

**Table 1:**
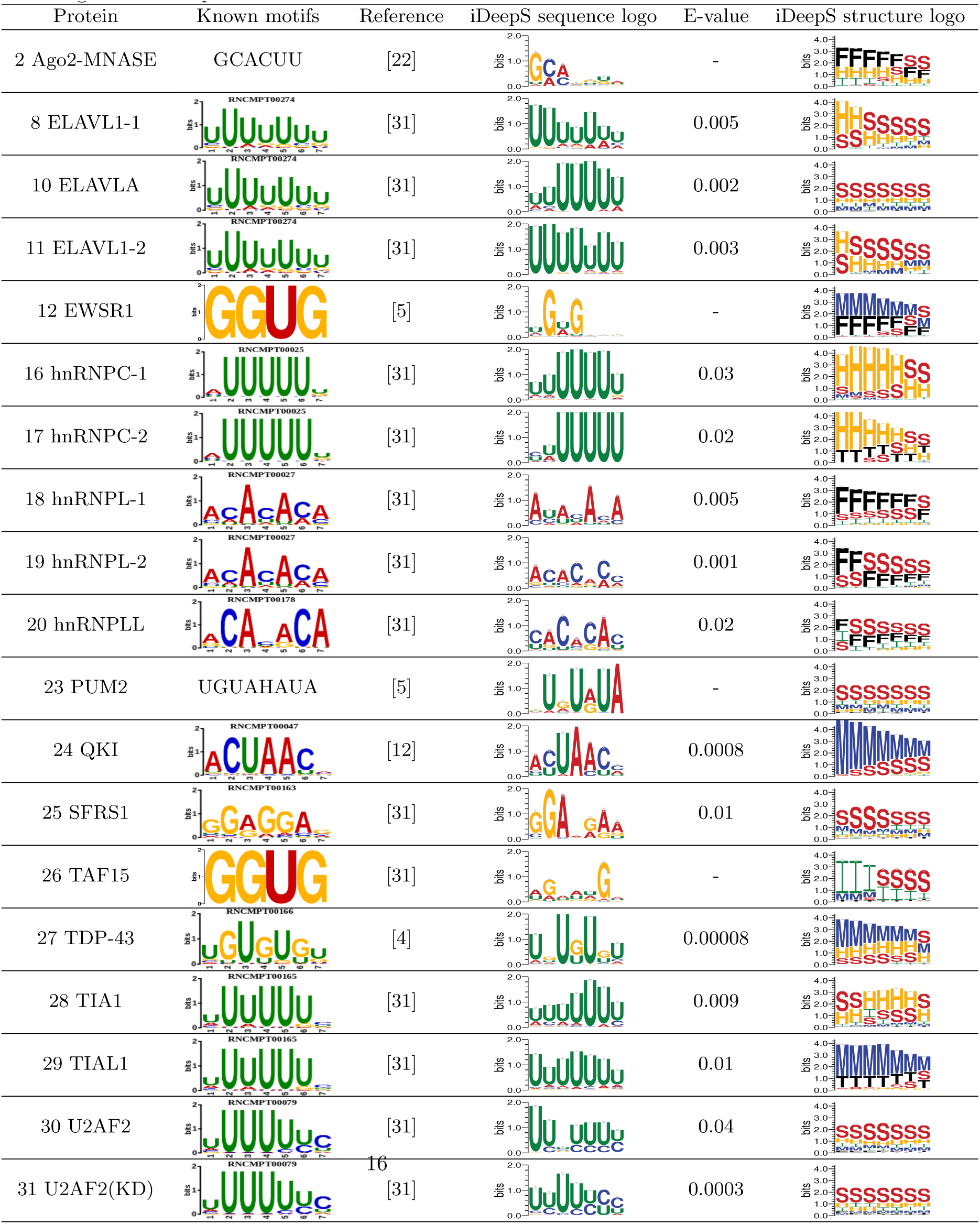
iDeepS captures known sequence motifs and structure motifs. The predicted sequence motifs are compared them against known motifs in study [31] from CISBP-RNA database and literatures. E-value is the expected number of false positives for the predicted motifs against known motifs using TOMTOM. The structure motifs are labelled as follows: stems (S), multiloops (M), hairpins (H), internal loops (I), dangling end (T) and dangling start (F). Note that these listed logos do not represent the full extent of the matched motifs.

Another advantage of iDeepS is that it enables discovery of structure motifs, which is not possible for iDeep [28]. iDeepS has demonstrated that most of proteins have preferences to generally structured regions. As shown in Table 1, the proteins in ELAVL protein family prefers to binding to stem-loop structure, which is consistent with the in vivo and in vitro binding data [11]. iDeepS also predicts that the protein U2AF2 prefers to binding to U-rich internal loop structures and hnRNPL proteins bind to external region with AC-rich, which agrees with the finding in [26].

Of particular interest, we further investigate the identified motifs for FUS, MOV10 and IGF2BP1-3 (Figure 3), who have no sequence motifs in CISBPRNA database. FUS has been found to bind to AU-rich stem-loop structure (Adjusted p-value: 1.55*e*^−2^ for structure motif) according to study [16], which is captured by iDeepS (Figure 3A). In addition, we also found the similar motifs to GraphProt for protein MOV10 with AU rich stem region (Figure 3 B, Adjusted p-value: 3.89*e*^−3^ for structure motif), and IGF2BP1-3 protein with CA dinucleotides multi-loop region (Figure 3C, Adjusted p-value: 5.01*e*^−5^ for structure motif). iDeepS discover another AC-rich stem-loop motif identified in [23] for Ago2 (Figure 3D, Adjusted p-value: 4.28*e*^−2^ for structure motif), which is different from the motif of Ago2 listed in Table 1. Compared to GraphProt, iDeepS is able to discover multiple binding sequence and structure motifs for each protein.

**Figure 3:**
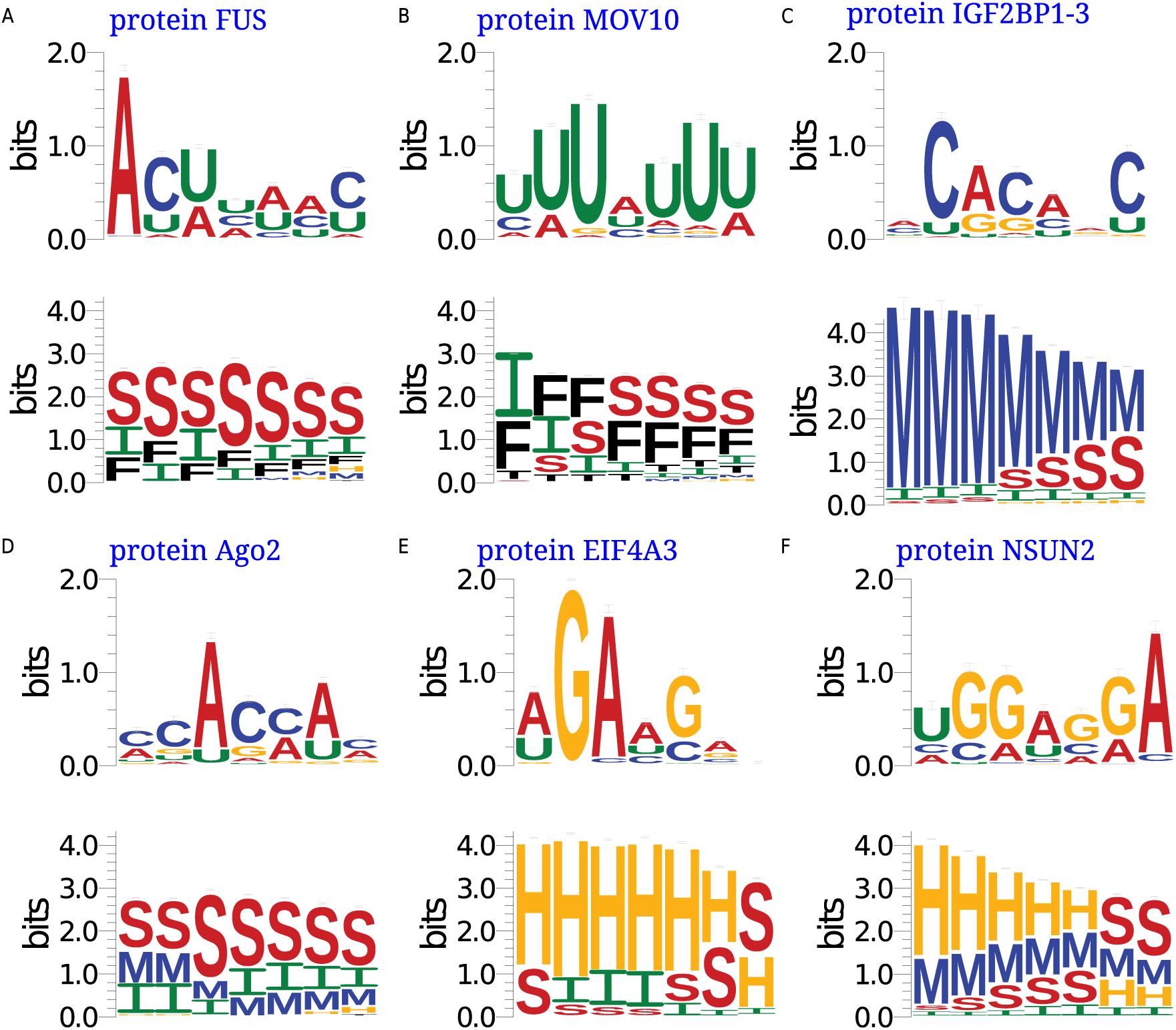
The identified novel binding sequence and structure motifs by iDeepS for RBPs. A. protein FUS. B. protein MOV10. C. protein IGF2BP1-3. D. protein Ago2. E. protein EIF4A3. F. protein NSUN2. In the structure motif logos, they are labelled as follows: stems (S), multiloops (M), hairpins (H), internal loops (I), dangling end (T) and dangling start (F)

We also discovered many novel motifs we could not verify against currently available knowledge. All discovered sequence and structure motifs by iDeep and and their enrichment score are available at https://github.com/xypan1232/iDeepS/tree/master/motif. For instance, iDeepS captures novel motifs for RBP EIF4A3 and NSUN2 (Figure 3E and F), their sequence motifs are enriched with adjusted p-value 5.18*e*^−53^ and 1.53*e*^−8^, respectively. Similarly, their structure motifs are enriched with adjusted p-value 4.20*e*^−3^ and 7.02*e*^−5^, respectively. They both show preference to harpin loop region. These discoveries have not been found by any studies, and need to verified in future studies.

### 3.3 Added value of BLSTM

To demonstrate the added value of BLSTM, we assess the iDeepS against a variant using only CNNs without the BLSTM layer. As shown in Figure 4, iDeepS yields better performance for most of 31 experiment. After taking the standard deviation of differences into consideration, iDeepS still significantly outperform the models only using CNNs on 10 experiments. The results indicate that BLSTM is able to capture better motifs for predicting RBP binding sites, which supports that the BLSTM can learn long-term dependencies.

**Figure 4:**
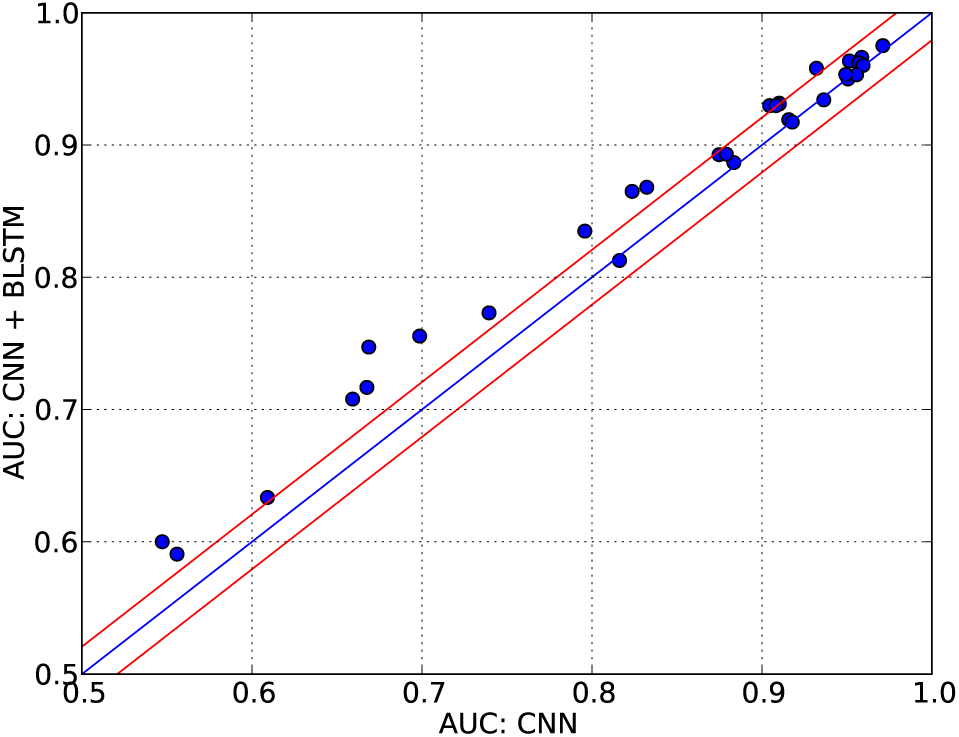
The difference of predictive performance using CNN + BLSTM and only CNN. On the y-axis the performance of the full model with CNNs and BLSTM is shown. The x-axis shows the performance of the model using only the CNNs without BLSTM. The two red lines indicate the standard deviation of the difference between only using CNN and using CNN + BLSTM.

DeepBind achieves an average AUC of 0.85 across 31 experiments by only using sequence CNN, which is a little better than 0.84 of simply concatenating the outputs from sequence and structure CNNs. The reason is that the structure information is predicted from sequences, leading to redundant information, which might hurt the model training. Both the model perform worse than iDeepS (AUC: 0.86) with BLSTM layer after sequence and structure CNNs. The results indicate BLSTM can learn long-term dependencies between sequence and structure motifs, which reduce the impact of redundant signals on training models for predicting RBP binding sites.

## 4 DISCUSSION AND CONCLUSIONS

In this study, we present a fully automatic deep learning based method iDeepS to infer both sequence and structure preferences of RBPs. The captured motifs align well with the previously reported binding motifs obtained from CISBP-RNA and literatures. Besides, iDeepS also discovers some novel motifs. The BLSTM layer in the iDeepS algorithm ascertains the long-term dependencies between sequence and structure motifs, which improves its predictive performance. We evaluated iDeepS on large-scale RBP binding sites derived from CLIP-seq datasets. iDeepS is able to predict the RBP binding sites on RNAs with higher accuracy than the state-of-the-art methods. Compared to existing black-box machine learning algorithms, iDeepS is able to find human-understandable sequence and structure binding motifs, which are expected to provide important clues for understanding the biological functional mechanisms of RNA and its binding protein RBP.

The training dataset in this study are originally downloaded from iONMF [35], in which the training positive binding sites are derived from CLIP-seq dataset, and the negative binding sites are constructed from genes that were not identified as interaction in any of 31 experiments. It has a stringent criteria to create the negative sites, even some genes may bind to certain RBPs but not bind to other RBPs. In this study, we use the same preprocessing to construct the training set for different RBPs. However, for some RBPs, iDeepS also fails in those cases where other existing tools also have low AUC values. The reason behind it is the quality of training dataset for this RBP might be low. Thus we need further improve the data quality with different strategies for different RBPs. In addition, different RBP families have different RNA-binding specificities, thus the training set is constructured per RBP.

The iDeepS not only can be used to predict RBP binding targets from RNA sequences, but also capture the binding preference. When there are available RNA sequences with potential target sites for RBPs of interest, then these sequences can be fed into iDeepS models. The iDeepS estimates the probability of those RNA sequences bound to certain RBPs. On the other hand, iDeepS also can identify the binding sequence and structure motifs. The captured sequence and structure context are an important basis for further research on their clinical impact. For example, these findings could contribute to discovering the mechanisms of diseases involving RBPs. Some structure specificities increase the possibility of the disruption of the structures within binding sites, which might cause diseases, e.g. protein FMR1 in fragile of X syndrome [9]. Furthermore, iDeepS has the potential application on predicting the effects of mutations. For example, we can mutate the nucleotides of binding sites, then use iDeepS to predict whether the new binding sites have a big shift compared to experimentally verified sites.

## 4.1 ACKNOWLEDGMENTS

This work was supported by the Natural Science Foundation of China (Nos.61671288, 31628003, 61462018, 61602176), Science and Technology Commission of Shanghai Municipality (No. 16JC1404300), China Postdoctoral Science Foundation Funded Project (No.2016M590337)

## 4.1.1 Conflict of interest statement

None declared.

